# Passive water exchange between multiple sites can explain why apparent exchange rate constants depend on ionic and osmotic conditions in gray matter

**DOI:** 10.1101/2025.05.27.655493

**Authors:** Nathan H. Williamson, Rea Ravin, Teddy X. Cai, Julian A. Rey, Peter J. Basser

**Affiliations:** Eunice Kennedy Shriver National Institute of Child Health and Human Development, National Institutes of Health, Bethesda, MD 20892, USA; Military Traumatic Brain Injury Initiative (MTBI^2^), Bethesda, MD 20814, USA; Uniformed Services University of the Health Sciences (USU), Bethesda, Maryland 20814, USA; The Henry M. Jackson Foundation for the Advancement of Military Medicine Inc. (HJF), Bethesda, Maryland 20817, USA; Celoptics, Rockville, MD 20850, USA; National Institute of General Medical Sciences, National Institutes of Health, Bethesda, MD 20892, USA

**Keywords:** Magnetic resonance in porous media, Single-sided NMR-MOUSE, Double diffusion encoding, NMR hydrophysiology, Active water cycling, Transcytolemmal water exchange, Water homeostasis, DEXSY, Sodium–potassium pump

## Abstract

Porous materials, such as biological tissue, often have heterogeneous microstructures where imbibed fluid experiences distinct environments on short timescales, but can exchange among different environments over long timescales. Nuclear magnetic resonance (NMR) methods such as diffusion exchange spectroscopy (DEXSY) can measure this exchange in water under steady-state and equilibrium conditions; however, modeling becomes more complex when more than two exchanging environments are involved. This complexity is particularly relevant in the central nervous system (CNS), where water diffusion and exchange at the cellular level play critical roles in homeostasis. While DEXSY can measure these processes, they may not be adequately modeled as two-site exchange between intracellular and extracellular spaces (ICS and ECS). Here we study the behavior of apparent exchange rate constants (AXR) estimated from DEXSY data numerically simulated using a three-site exchange model (3XM). The 3XM is based on gray matter microstructural characteristics, incorporating both transmembrane exchange between ECS and ICS and geometric exchange between environments within ICS where water mobility differs due to the complex architecture of neurons, glial cells, and the ECS. Inspired by the Na^+^/K^+^–ATPase pump–leak model of cell volume maintenance, the 3XM accounts for effects of osmolytes, ions, and voltage on ECS and ICS volume fraction. The model predicts a significant reduction in AXR and a smaller decrease in apparent diffusion coefficients (ADC) following the level of membrane depolarization expected from Na^+^/K^+^–ATPase inhibition. These changes were reversed by the addition of membrane-impermeable ECS osmolytes, independent of voltage, in agreement with previous experiments. While the exchange rate constants for each pathway simply follow first-order kinetics, the AXR’s sensitivity to these pathways depends on the ECS volume fraction. When ECS is present, transmembrane exchange dominates, but when cells swell following pump inhibition, geometric exchange becomes the dominant pathway.

## 1. Introduction

Soft matter and porous media physics is often concerned with understanding the migration of molecules between sites under steady-state conditions [1–3]. These “exchange” processes are crucial in various applications, ranging from catalysis [4, 5] and separations [6, 7] to cell biology [8, 9] and medicine [10, 11]. In biological tissues, exchange of water across cell membranes is a physiologically important characteristic of homeostasis [12]. However, few techniques can measure exchange, and those that do often require complicated experimental setups and operate under specific constraints [13–15]. For instance coherent anti-Stokes Raman scattering (CARS) microscopy has been used to visualize water permeability of the arterial wall, but this highly specialized method involves D_2_O tracer imaging in a superficial tissue layer [16]. CARS cannot measure exchange deeper in tissue or on timescales faster than the media can be washed between H_2_O and D_2_O.

NMR can also measure steady-state fluid transport, but has a different set of advantages and limitations defining its niche. NMR works by encoding and detecting the resonance of nuclear spins on mobile molecules, such as protons (^1^H) on water (H_2_O), within an external magnetic field [17, 18]. It may be the only method capable of noninvasively measuring exchange of endogenous fluid molecules deep within optically turbid materials on millisecond to second timescales. In living tissue, the apparent (time-dependent) self-diffusion coefficients (ADC) of water are on the order of 1 µm^2^/ms. This enables the study of water exchange between sub-micron to micron-scale cellular compartments, averaged over a much larger imaging voxel or active region [11, 19–23]. However, measuring exchange dynamics is an inverse problem that requires modeling.

Modeling exchange in central nervous system (CNS) tissue is crucial for the diffusion MRI community [11, 21, 23–28]. Although exchange is often considered slow enough to be neglected in white matter models [29], recent studies have shown it to be much faster and essential for accurate modeling of gray matter [20–22, 28, 30–33]. Yet a two-site exchange model (2XM) involving transmembrane exchange between a homogeneous intracellular space (ICS) and extracellular space (ECS) is typically assumed. Diffusion MR studies have only begun to address the impacts of tissue heterogeneity[34–42]. This poses challenges and opportunities, as accurate modeling and understanding could provide new imaging biomarkers.

Transmembrane water exchange is typically assumed to be a passive process driven by thermal motion, resulting in a rate constant for the turnover of intracellular water that is proportional to membrane diffusive permeability and surface-to-volume ratio (SVR) [43, 44]. Some studies suggest an additional component of water exchange linked to active transport [45–54]. The Na^+^/K^+^–ATPase is the primary active transporter in animal cells [55]. It utilizes the energy from one ATP phosphate bond to transport three Na^+^ out of the cell and two K^+^ into the cell. Per unit mass, the CNS is the most metabolically active organ in the body, and continual Na^+^/K^+^–ATPase activity accounts for about half of that energy [56]. The resulting ionic gradients facilitate secondary active and passive ion transport, which play a role in both functional processes, such as neuronal firing [57, 58], and homeostatic processes, like maintaining and regulating cell volume [59–62]. Studies have suggested that water may also be transported or cycled with ions, and that transmembrane water exchange could serve as a biomarker for metabolism [45–54].

Much of the evidence for “active water cycling” comes from the strong inhibition of active transport on water exchange using various channel blockers [45, 48, 50]. For instance, Bai *et al.* found that inhibiting Na^+^/K^+^–ATPase activity in organotypic culture slices with the drug ouabain reduced the exchange rate constant by 45% [48]. Similarly, in our *ex vivo* neonatal mouse spinal cord studies, we observed that ouabain reduced the apparent exchange rate constant (AXR) by 70% from 140 s^−1^ to 40 s^−1^ [53]. (While in previous studies we used *k* for the estimated apparent exchange rate constant, here we use AXR to avoid confusion with the ground-truth exchange rate constant *k* defined for the 2XM.) Although both studies reported cellular swelling, the extent of SVR reduction was considered insufficient to explain the drop in AXR. However, more recent findings revealed that osmolytes rescued the AXR, even when Na^+^/K^+^–ATPase activity remained inhibited by ouabain [38]. These results, along with other experiments, led us to reject the hypothesis that AXR is linked to active water cycling in the neonatal mouse spinal cord. Instead, results suggest that AXR is related to osmotic conditions, which are maintained by cellular homeostasis or controlled by the bathing media [38]. The goal of this study is to develop a minimal model that qualitatively reproduces the dependence on osmotic swelling and shrinking observed in that companion study [38]. Our hypothesis is that AXR behavior can be explained by changes in volume fractions due to osmotic swelling and shrinking in a multisite exchange model involving only passive exchange.

In this paper we first present the theoretical background of the diffusion exchange spectroscopy (DEXSY) experiment and how it is performed using the static gradient of a fringe field or low-field, high-gradient system. We also cover the theory of exchange modeling as well as basic aspects of cell volume maintenance. The latter motivates the use the pump–leak model (PLM) [62] pertinent to predicting the dependence of site volume fractions on osmotic swelling and shrinking. We then describe the PLM, 2XM and three-site exchange model (3XM) simulation methods. Finally, we present simulation results for the PLM, 2XM and 3XM along with extended simulations. We qualitatively compare model predictions of how AXR and ADC depend on osmotic conditions to experimental results from a separate study on the neonatal mouse spinal cord [38]. At this stage, we do not fit the 3XM to data. However, since the neonatal mouse spinal cord consists primarily of gray matter [63, 64], this model may be relevant to future developments of gray matter diffusion MRI models which are aimed at estimating tissue parameters.

### 2. Theory

### 2.1 SGSE DEXSY and the diffusion exchange ratio (DEXR) method

Whereas diffusion encoding is more commonly performed with pulsed gradient spin echos (PGSE) [65] we use static gradient spin echoes (SGSE). With SGSE, gradient and spin echoes are formed by using hard RF pulses to modulate the effect of a static gradient from a strongly decaying *B*_0_ field [66]. With permanent single-sided magnets such as the NMR MOUSE [67], these static gradients can be greater than 10 T/m, for instance *g* = 15.3 T/m in our companion study [38]. While diffusion encoding with PGSE typically involves varying the gradient amplitude [65], with SGSE it involves varying τ (= 1/2 the echo time) to achieve the desired *b* value, where

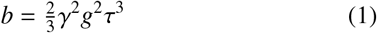

and γ is the gyromagnetic ratio [68]. While magnetization from freely diffusing spins far from surfaces, and spins that have completely coarse-grained (averaged) over interactions on shorter timescales [69] attenuate proportional to exp(−*b*ADC), the magnetization from water for which the extent of diffusion in the gradient direction is bounded by surfaces attenuates much slower.

The distinguishing feature is the structural length scale 𝓁_*s*_ between surfaces surrounding the water relative to the dephasing length scale, 𝓁_*g*_ = (*D*_0_/γ*g*)^1/3^ and diffusion length scale 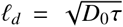 [70]. Assuming a water self-diffusion coefficient *D*_0_ = 2.15 µm^2^/ms at 25°C and a strong static *g* = 15.3 T/m, 𝓁_*g*_ = 0.8 µm [20, 71, 72]. Moreover, *l*_*g*_ changes weakly with *g* due to the 1/3 scaling. Clinical high-field PGSE with *g*_*max*_ = 0.3 T/m can still access 𝓁_*g*_ = 3 µm [73], and a similar framework should apply. 𝓁_*g*_ is roughly the distance over which diffusing spins dephase by 2π radians. It can be considered the microscale “resolution” of SGSE diffusion encodings (or when gradient duration equals the time between the leading edges of the gradient pulses, i.e., δ = Δ, with PGSE). Water will feel the confining effects of membranes when 𝓁_*d*_ is greater than or similar to 𝓁_*s*_ and 𝓁_*g*_. In this regime, 𝓁_*s*_ < 𝓁_*g*_ results in motional averaging of signal within the compartment [74], whereas 𝓁_*g*_ < 𝓁_*s*_ results in localization of signal within ~*l*_*g*_ of bounding surfaces oriented perpendicular to the gradient direction [70, 72, 73, 75–79]. The idea of compartments breaks down in the localization regime and hence we ignore it. For our purposes, SGSE diffusion encoding distinguishes compartments or sites based on the local 𝓁_*s*_ in the direction of *g* relative to 𝓁_*g*_ and 𝓁_*d*_.

The most common approach to measuring exchange is to vary the diffusion time (referred to as τ above), using PGSE or pulsed-gradient stimulated echo (PGSTE) diffusion experiments [80]. However, these methods are not specific because factors other than exchange affect the signal attenuation in this single time dimension [31, 32]. 2D-exchange NMR methods such as diffusion exchange spectroscopy (DEXSY) [81] (also referred to as double diffusion encoding [82]) reduce model degeneracy by separating the encoding of site-specific diffusion from the time period during which mixing or exchange between sites is observed (i.e., the mixing time, *t*_m_) [83, 84]. Note that *t*_m_ stores signal formed at a spin and gradient echo and is different from a stimulated echo. Another distinguishing feature is that the resulting DEXSY data can be converted into 2D exchange distributions using a 2D inverse Laplace transform [85, 86]. However, this requires acquiring enough points in the encoding space to obtain a stable and resolved distribution, which often takes too long for biological applications [87, 88]. Additionally, the inversion process assumes that a Gaussian kernel can accurately describe the relationship between encoding and signal attenuation. However, when the kernel fails to correctly model this relationship, it leads to artificial exchange components. This issue commonly arises in DEXSY studies of porous media, where environments exhibit non-Gaussian restricted diffusion [71].

To overcome these challenges, we developed and validated a DEXSY-based NMR and MRI method called diffusion exchange ratio (DEXR) MR for rapidly measuring an apparent exchange rate constant AXR in as few as three *t*_m_ values, with one or two encoding combinations per *t*_m_ (the second point accounting for *T*_1_ relaxation during *t*_m_) [83, 89, 90]. DEXR estimates of AXR are valid for non-Gaussian diffusion, as such effects influence signals equally across all *t*_m_ values [71, 90]. However, like the DEXSY-based methods, filter exchange spectroscopy (FEXSY) and filter exchange imaging (FEXI) [91–93], this approach assumes that only two components are exchanging.

Here we explain DEXR for the SGSE DEXSY, although the concept is the same for DEXSY performed using pulsed gradients. The DEXSY pulse sequence involves two diffusion encodings separated by *t*_m_ (Fig. 1 A).

**Fig. 1:**
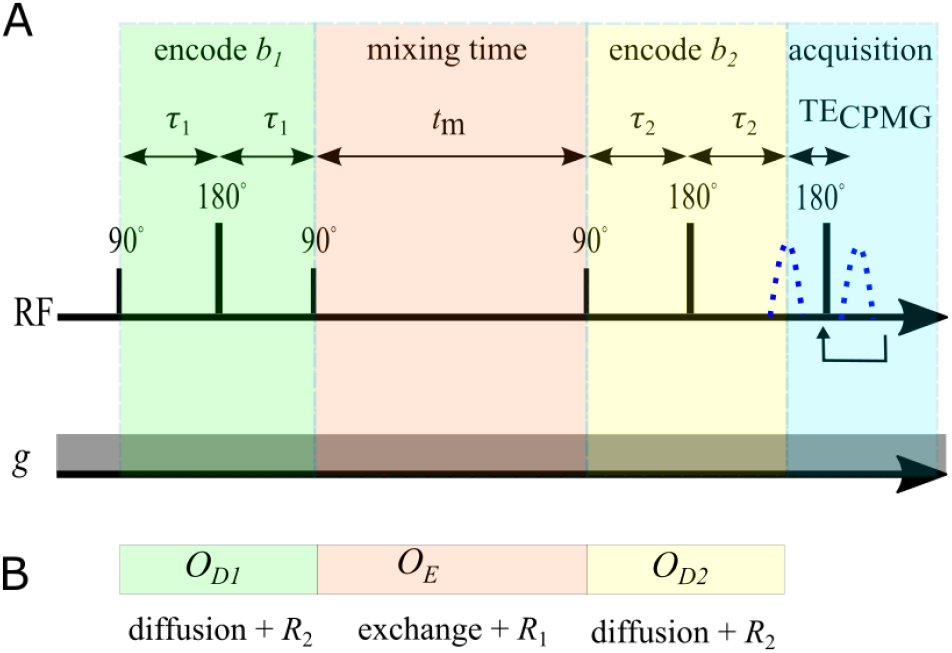
A) Encoding portion of the SGSE DEXSY pulse sequence, showing radiofrequency (RF) pulses and the first acquired echo signal (blue dotted line). In practice, signal can be refocused in a CPMG train to boost SNR (see [83]). The static gradient *g* is always on. Diffusion encoding times τ_1_ and τ_2_ are varied to set *b*_1_ and *b*_2_. B) Operators used to simulate SGSE DEXSY signals.

The basic idea behind DEXR is quite simple. First, we recognize that *t*_m_ is the key dimension containing exchange information [90]. For maximum exchange contrast, *b*_1_ and *b*_2_ values are set equal and optimized to dephase more mobile spins but refocus less mobile spins. During the first diffusion encoding block, spins in more mobile compartments dephase. During *t*_m_, spins are able to migrate to new compartments without additional dephasing. In the slow exchange regime (*t*_m_≪1/*k*) very little exchange occurs, there is not much additional dephasing during the second diffusion encoding block, and the echo signal is simply attenuated by *b*_*s*_ = *b*_1_ + *b*_2_. As *t*_m_ is increased into the intermediate exchange regime (*t*_m_~1/*k*) and towards the fast exchange regime (*t*_m_≫1/*k*), the more mobile compartments are replenished with spins from the less mobile compartments and the second diffusion encoding block causes additional attenuation. This additional attenuation is exchange contrast.

However, *T*_1_ relaxation also occurs during *t*_m_. This can be accounted for by normalizing the signal from each mixing time by a point with *b*_1_ and *b*_2_ near zero, but can lead to biases because the ensemble average *T*_1_ can be different from compartment *T*_1_ values. Instead, we can realize that setting one diffusion encoding block (*b*_1_ or *b*_2_) near zero and varying the other is like performing a diffusion–*T*_1_ correlation experiment and does not provide exchange contrast [83]. A signal acquired with *b*_1_ = 0 and *b*_2_ = *b*_*s*_ results in the same level of (Gaussian) diffusive attenuation and *T*_1_ attenuation as the point with *b*_1_ = *b*_2_ = *b*_*s*_/2. Normalizing by this point isolates attenuation due to exchange and results in the “DEXR signal”.

In practice, two diffusion encoding blocks with *b*_*s*_/2 and *t*_m_ = 0 can result in more attenuation than one encoding block with *b*_*s*_ due to non-Gaussian diffusion [71, 94] (e.g. localization and motional averaging [72]), but this additional attenuation is the same across mixing times and can be lumped into a model parameter related to the non-Gaussian signal fraction [71, 90].

To fully characterize the exchange process, DEXR signals should be acquired at multiple *t*_m_ values spanning the slow, intermediate, and fast exchange regimes.

### 2.2 Multisite exchange signal modeling

Multisite steady-state exchange can be modeled by formulating the Bloch equations as a system of matrix differential equations [35, 95–100]. This effectively involves multiplication of matrix exponentials, signified by expm(). However, it assumes that magnetization in each individual site is well mixed. This is called the “fast diffusion” regime in relaxation [101] and is a condition for “barrier-limited exchange” in diffusion [27]. It also assumes that signal relaxation/attenuation in each site can be modeled as an exponential decay. Lastly, it assumes exchange can be modeled with first-order kinetics, meaning that the unidirectional flux from one site to another, e.g. from site *i* to site *j*, is equal to the number of spins occupying site *i* multiplied by a rate constant *k*_*i j*_ where the first and second subscript indices signify where the spins are coming from and going to.

For *N*-site exchange in DEXSY, the initial (*t* = 0) normalized signal is **S_0_** = [*f*_1_, …, *f*_*N*_]^′^ where ^′^ signifies the transpose operator and *f*_1_ … *f*_*N*_ are signal fractions for sites 1 through *N* and sum to 1.

The effect of diffusion is modeled as **S** = expm(−*b*_1,2_**D**)**S_0_** with the diffusion matrix

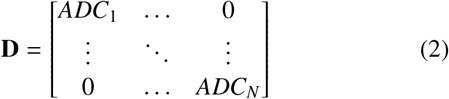

Similarly, spin–spin relaxation during the periods spins spend in the transverse plane, and spin–lattice relaxation during the magnetization storage period, can be modeled as **S** = expm(−*t***R**)**S_0_** with relaxation matrices **R**_2_ or **R**_1_ with diagonals containing *R*_2_ or *R*_1_ relaxation rate constants for each site.

The signal evolution due to exchange alone is given by

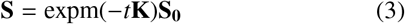

where

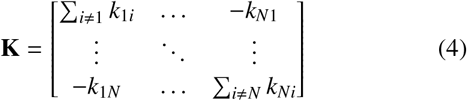

is the exchange matrix. **K** has the effect of mixing the magnetization of signal components, but conserves the total magnetization. This means that the total balance of magnetization leaving and coming into each site is preserved

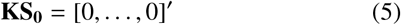

and requires that each column of **K** sum to 0 [102]. At equilibrium, the forward and backward flux between any two sites is also balanced

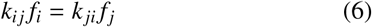

This “principle of detailed balance” [96], is not necessarily true in multisite exchange systems under nonequilibrium conditions. Some studies report violation of Eq. 6 due to circular exchange between sites, e.g., from site 1→2→3→1 [103, 104]. However total magnetization is still conserved (Eq. 5 still holds) under steady-state conditions with no net flow.

These matrix exponentials also enter into operators for each of the encoding periods of the SGSE DEXSY sequence (see Fig. 1 B). The operators for the first and second diffusion encoding blocks are *O*_*D*1,2_ = expm(−*b*_1,2_**D−**2τ_1,2_**R**_2_ −2τ_1,2_**K**). The operator for the mixing time is *O*_*E*_ = expm(−*t*_m_(**K** + **R**_1_)). This provides a convenient way to simulate SGSE DEXSY signals

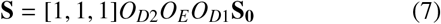

Dortch *et al.* provide detailed derivations of the analytical solutions to the signal behavior generalized for *N*-site exchange [96]. An important insight is that the measured exchange rate constants are the nonzero eigenvalues of **K**+**R**_1_. In the case that relaxation can be compensated, such as in the DEXR approach, the measured AXRs are the nonzero eigenvalues of **K**.

### 2.3. Two-site exchange model

NMR and MRI studies often assume a two-site exchange model (2XM). These sites or environments are distinguished by signal decay or attenuation (effectively having distinct ADC values). Spins completely sample these environments faster than they exchange between them. As an important example, the plasma membrane separating the ICS and ECS of cells also acts as a barrier to diffusion, simultaneously restricting diffusion in the ICS on short timescales but permitting exchange on longer timescales. Water in the ECS may appear more mobile because it is a connected space, albeit a narrow and tortuous one. When the timescale to diffuse across the cell is shorter than the timescale to exchange, this is called barrier-limited exchange and the behavior is described by a 2XM [27].

Simulation of DEXSY signal from two-site exchange between sites *a* and *b* with fractions *f*_*a*_ and *f*_*b*_ can follow the same approach outlined above. In this case,

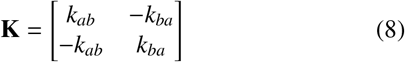

with eigenvalues

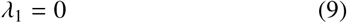

due to the total balance (Eq. 5) and

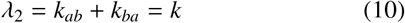

which defines the ground-truth exchange rate constant *k*. Using this, *f*_*b*_ = 1− *f*_*a*_, and Eq. 6 allows for the exchange matrix to be expressed as [98]

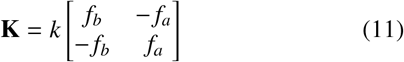

### 2.4. Three-site exchange model for gray matter

Some heterogeneous systems including biological tissues involve molecules exchanging between multiple sites and may be better described by a multisite exchange model. For instance, using *T*_2_–*T*_2_ or relaxation exchange spectroscopy (REXSY), Dortch *et al.* provided evidence of three-site exchange from water moving between the ECS, myelin sheaths, and intra-axonal space in CNS white mater [105]. Diffusion in gray matter may also show multisite exchange, but for different reasons. DEXSY distinguishes exchange between compartments based on water translational mobility in the direction of the magnetic field gradient *g*, not based on water’s location in the ECS or ICS *per se* [106]. Gray matter microstructure is characterized by the presence of branching cellular processes with sub-micron radii, and cell bodies (soma) with radii ranging from a few microns to tens of microns [107]. Consistent with this picture, in the neonatal mouse spinal cord, fluorescent images of dye-loaded motoneurons and interneurons [108] and of selectively labeled astrocytes [109] show cell soma with processes branching in every direction. Water diffusion within sub-micron processes oriented perpendicular to the gradient direction may appear restricted. Water diffusing in larger soma and in processes running parallel to *g* may appear more mobile. This raises the possibility of geometric exchange between intracellular regions that exhibit different mobilities along the gradient direction. While various geometric exchange pathways could exist, potential intracellular sources include diffusion along bending [110] or branching processes [106], between cell soma and processes [111, 112], or between spines and shaft of dendrites [42, 113]. Transmembrane exchange also occurs between neuronal and glial intracellular spaces (ICS) and the ECS [30].

This motivates a three-site exchange model (3XM) for diffusion in gray matter with exchange between an ECS compartment (*a*), a more mobile ICS compartment (*b*), and a less mobile ICS compartment (*c*), depicted in Fig. 2. The model proceeds in the same fashion outlined for multisite exchange modeling and makes the same assumptions.

**Fig. 2:**
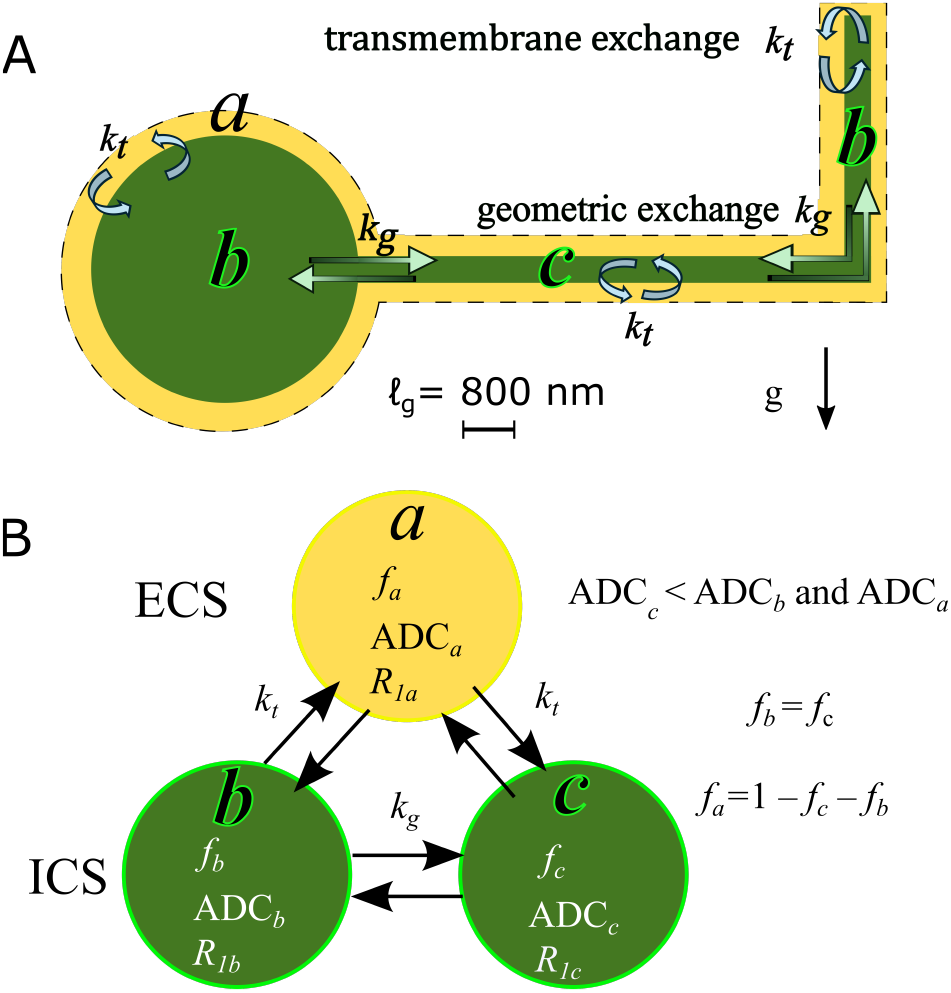
A) Three-site exchange model (3XM) for gray matter with the ECS (compartment *a*), cell bodies and processes oriented parallel to *g* (*b*), and cellular processes oriented perpendicular to *g* (*c*). ADC_*c*_ is lower than ADC_*b*_and ADC_*a*_ because 𝓁_*s*_ ≲𝓁_*g*_. Water in compartments *b* and *a* exhibits greater mobility along *g* since membrane length scales in that dimension exceed 𝓁_*g*_.Transmembrane exchange *k*_*t*_ occurs between the ECS and ICS compartments:*a*–*b* and *a*–*c*. Geometric exchange *k*_*g*_ occurs between ICS compartments: *b*–*c*.B) Relationships between compartments.

The exchange matrix is

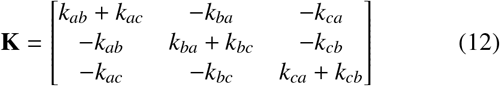

In analogy to the relationship between Eqs. 8 and 11, we define

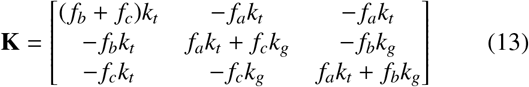

where *k*_*t*_ and *k*_*g*_ are the transmembrane and geometric exchange rate constants, respectively. Eq. 13 satisfies the total balance (Eq. 5) and all detailed balances (Eq. 6). The eigenvalues of Eq. 13 are

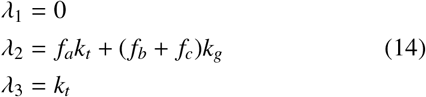

Where λ_1_ = 0 is a result of the total balance (Eq. 5), λ_2_ is the “spectral gap” or slowest relaxing mode, and λ_3_ is the fastest relaxing mode. Below, we will use a 2XM-based fit to estimate AXRs from data simulated with three sites. To gain some intuition for what AXR should be, we can look at the extremes. When *f*_*a*_ = 0, the system is expected to reduce to two-site exchange between sites *b* and *c* and AXR = *k*_*g*_. As *f*_*b*_ + *f*_*c*_ approach 0, geometric exchange between sites *b* and *c* is expected to become less apparent and AXR = *k*_*t*_. These conditions are fulfilled if the AXR is a sum of λ_2_ weighted by *f*_*b*_ + *f*_*c*_ and λ_3_ weighted by *f*_*a*_.

### 2.5. Cell volume maintenance

This section explains why the cell volume and hence the ICS and ECS volume fractions (*f*_o_ and *f*_i_) are intimately related to Na^+^/K^+^–ATPase activity under normal conditions. This will be important for defining the compartment fractions (*f*_*a*_, *f*_*b*_, *f*_*c*_) for the 2XM and 3XM simulations.

All cells have plasma membranes which are semipermeable to water and ions but prevent metabolites, proteins, and nucleic acids from permeating out. These trapped ICS impermeants, with total moles *x*_i_, carry a net-negative charge *Z*, which is balanced by intracellular ions to maintain electroneutrality:

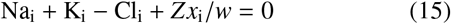

where *w* is the cell volume. Following Kay (2017) [62], we include only the most prominent monovalent ions for simplicity, while acknowledging that other ions such as Ca^2+^, Mg^2+^ and HCO^−^_3_ are present in the media and play important roles [61, 114]. The impermeants and associated cations exert an osmotic pressure π_i_ on the membrane. The van’t Hoff equation for ideal solutions can be used to estimate π for a given solute concentration *c* and absolute temperature *T* [115],

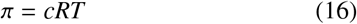

where *R* is the ideal gas constant. While the ICS environment deviates significantly from ideality, Eq. 16 can be used to provide a rough estimate of the pressure contribution from soluble components. With the concentration of intracellular impermeants and associated cations being on the order of 10 mM, π_i_ is predicted to be on the order of 100 kPa [116]. Plants, fungi, and most bacteria evolved rigid cell walls capable of counteracting this pressure. Animal cells, in contrast, lack cell walls because they need to be distensible to facilitate movement. Additionally, parenchyma tissue (the functional non-epithelial tissue within organs) typically has an ECS.

If hydrostatic pressures are considered negligible, the intracellular osmotic pressure must be balanced by an extracellular osmotic pressure π_i_ = π_o_ for a stable volume to exist. By Eq. 16, this requires that intracellular and extracellular osmolarities be equal:

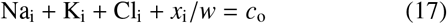

Here, *c*_o_ = Na_o_ + K_o_ + Cl_o_ + *s*_o_ represent ECS concentrations, typically assumed to match the surrounding fluid. This assumption holds for isolated cells but may not in tissues. *s*_o_ accounts for uncharged osmolytes, which could be endogenous or added to the bathing media as was done in the *ex vivo* experiments we will compare our simulations to [38]. Subtracting Eq. 15 from Eq. 17 and rearranging results in an equation for cell volume [62]

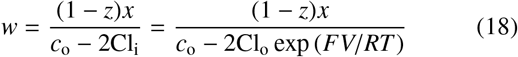

where Cl_i_ = Cl_o_ exp (*FV*/*RT*) is the Nernst equation for chloride’s electrochemical potential with Faraday constant *F* and voltage *V*. For later reference, at *V* = −48 mV and −10 mV (and *T* = 298 K), the fraction of chloride partitioning is predicted to be Cl_i_/Cl_o_ = exp (*FV*/*RT*) = 0.15 and 0.68, respectively. This shows that volume and voltage are interconnected normally and that cells maintain volume by partitioning chloride.

The non-equilibrium state is maintained by active ion transport by the Na^+^/K^+^–ATPase, coupled with higher permeability of K^+^ relative to Na^+^. K^+^ flows or “leaks” passively down its electrochemical potential gradient and brings Cl^−^ with it to maintain electroneutrality. This, in turn, drives water movement to equilibrate osmotic pressure. This is called the “pump–leak model” [59, 62]. Notably, the so-called leaking ions are actually involved in cellular functions not accounted for in this model.

## 3. Methods

### 3.1. Pump–leak model

The pump–leak model (PLM) of cell volume maintenance provided by Kay (2017) was used to predict how perturbations affect cell volume *w* and transmembrane voltage *V* [62].The PLM approximates intracellular ion concentration and *w* changes over time from the active flux of Na^+^ and K^+^ by a defined Na^+^/K^+^–ATPase pump rate and passive fluxes of Na^+^, K^+^, and Cl^−^ based on their electrochemical potential. Fluxes are approximated using the finite difference method.

The model assumes electroneutrality (Eq. 15) and no transmembrane osmolarity gradient (Eq. 17). While electroneutrality is a reasonable assumption, the absence of an osmolarity gradient may not hold in extreme cases, such as swollen or shrunken cells, particularly in tissue where mechanical pressures likely play a significant role. Water permeability is assumed to be much higher than ion permeability and is not modeled directly. Instead, the cell volume changes so that the intracellular and extracellular osmolarities are equal at the end of each timestep.

The net charge of the intracellular impermeants is *z* = −1. Voltage is modeled based on the net charge of the intracellular ions and the (constant) capacitance of the membrane. The normal media condition was defined as Na_o_ = 128 mM, K_o_ = 4 mM, and Cl_o_ = Na_o_ + K_o_ = 132 mM. This is similar to the artificial cerebrospinal fluid (aCSF) media composition used for *ex vivo* experiments [38, 53] except that it omits divalent cations, sodium bicarbonate, and glucose. Other parameter values were the same as in Kay (2017) and can be found there [62]. Volumes and voltages were taken as the values obtained at the final timestep. This time was sufficiently long for systems to reach steady-state (if there existed a stable steady-state), as determined by volume and voltage not changing when total time was increased by a factor of 10.

For all “PUMP ON” conditions, the Na^+^/K^+^–ATPase pump rate was set to the value which maintained a transmembrane voltage of *V* = −48 mV in the normal (128 mM NaCl) media. This voltage is based on intracellular recordings from motoneurons in the *ex vivo* neonatal mouse spinal cord (−48.4 ±5 mV) [117]. Since this value was recorded from a limited number of neuronal (and not glial) cells, it may not fully represent all CNS cells. Note also that in reality there are multiple Na^+^/K^+^– ATPase isoforms and their rates can vary [118–120].

For “PUMP OFF” conditions, the pump rate was set to zero. Under this condition in normal media, the PLM predicts unchecked cell swelling and a gradual voltage drop toward zero without reaching steady state, due to osmotic imbalance from intracellular impermeants. However, experimental observations suggest a limit to cell swelling [38].

### 3.2. Prediction of f_o_ from osmotic balance

The first step in both the 2XM and 3XM was to predict the ECS volume fraction (*f*_o_) for a specified osmolarity and voltage. To do this we first utilized prior knowledge that cells in tissue do not swell infinitely and hence there must be an extracellular pressure that builds up and limits the extent of cell swelling. While we do not know the source of this pressure, as a placeholder to model the behavior in a simple way, we fixed the total volume of ECS and ICS to *w*_tot_ and added uncharged impermeants trapped in the ECS (unable to permeate to the bath or across the cell membrane) with total moles *x*_o_. The fixed *w*_tot_ limits the maximum extent of cell swelling. It could conceivably account for the dura surrounding the CNS which holds the tissue together. However, it neglects the distensibility of the dura which could potentially allow *w*_tot_ to vary. *x*_o_ was arbitrarily set to 1/50^th^ of *x*_i_. Regardless of how minuscule *x*_o_ is, it forces *f*_o_ to be greater than zero and to vary smoothly because the associated osmolarity builds up asymptotically with *x*_o_/(*f*_o_*w*_tot_) as *f*_o_ decreases towards zero. It could conceivably model the effect of extracellular matrix, however it neglects the stiffness and charge of extracellular matrix (ECM) which cause osmotic pressure to increase greater than linearly with concentration and to depend on the ionic strength of the ECS solution [121].

Eq. 18 was re-derived with the additional *x*_o_ and *w*_tot_ terms and re-arranged, leading to an analytical equation to predict *f*_o_ as a function of *s*_o_ and Cl_o_ concentrations and *V* under steadystate conditions:

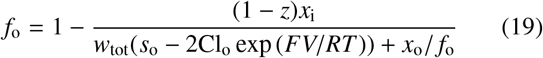

Since *f*_o_ is on the RHS as well, its value is found iteratively. In this study, *T* = 298 K, Cl_o_ = 132 mM, and *s*_o_ was varied.

Note that unlike in the PLM where voltage was predicted for a given pump rate, here *V* is specified. To model conditions where the Na^+^/K^+^–ATPase is functioning normally, we used *V* = −48 mV, based on intracellular recordings [117] discussed above. To model conditions where the Na^+^/K^+^–ATPase is inhibited, we set the voltage to *V* = −10 mV, reflecting recordings under hypoxic or terminal depolarization conditions in various CNS tissue models, which are thought to represent the end result of Na^+^/K^+^–ATPase inhibition [122].

*w*_tot_ was kept constant but needed to be defined at the start. To do this, first *w* was predicted using Eq. 18 with normal media and *V* = −48 mV. Then *w*_tot_ was defined by setting an initial *f*_o_. Here, *f*_o_ = 0.3 was chosen based on real-time iontophoresis with tetramethylammonium measurements of 0.27 in the P8– P10 mouse spinal cord slice model and the trend towards larger ECS fraction in younger animals [123, 124].

### 3.3. Numerical simulations of two- and three-site exchange

With *f*_*a*_ = *f*_o_ defined from Eq. 19, the intracellular compartment fraction(s) was/were constrained: *f*_*b*_ = 1− *f*_*a*_ in the case of the 2XM or *f*_*b*_ + *f*_*c*_ = 1 −*f*_*a*_ in the case of the 3XM.

DEXSY data was numerically simulated using the 2XM described in section 2.3 for comparison to 3XM results. The ADC and *R*_1_ values were set to ADC_*a*_ = 1 µm^2^/ms, ADC_*b*_ = 0.1 µm^2^/ms, and *R*_1*a*_ = *R*_1*b*_ = 1 s^−1^. In Eq. 11, *k* was set to 300 s^−1^, the same value used for *k*_*t*_ in the 3XM.

Numerical simulations were performed using the 3XM described in section 2.4 under several conditions. For simplicity,*f*_*b*_ and *f*_*c*_ were set equal, *f*_*b*_ = *f*_*c*_ = (1 −*f*_*a*_)/2. ADC values for the compartments were defined as ADC_*a*_ = 1 or 1.7 *f*_o_ µm^2^/ms, ADC_*b*_ = 1, 0.5, or 1.5 µm^2^/ms, and ADC_*c*_ = 0.1 µm^2^/ms, depending on the simulation (defined in Figure captions). These values were chosen in an effort to recapitulate experimental results recorded on the *ex vivo* spinal cord at 25°C [38] and in consideration of *in vivo* brain diffusion MRI literature (although knowingly different due to the temperature being 37°C and the diffusion encoding times being longer) [29, 125, 126].

The process for numerically simulating DEXSY signals was similar for the 2XM and 3XM. We describe the process here for the 3XM. DEXSY signals were simulated using Eq. 7 and the operator formalism presented in sections 2.2 and 2.4. Matrix exponentials were calculated using the function expm() in MATLAB 2024a. Equilibrium magnetization was **S_0_** =[*f*_*a*_, *f*_*b*_, *f*_*c*_]^′^. The operators for the first and second diffusion encoding blocks were *O*_*D*1,2_ = expm(−*b*_1,2_**D**) and used the diffusion matrix defined by ADC_*a*_, ADC_*b*_, and ADC_*c*_. We assumed no exchange and no spin–spin relaxation during encoding based on τ ≪1/*k* and τ ≪*T*_2_, although they can be included [96, 98]. The exchange operator was *O*_*E*_ = expm(*t*_m_(**K** + **R**)) and used the exchange matrix shown in Eq. 13 and spin–lattice relaxation matrix defined by *R*_1*a*_ = *R*_1*b*_ = *R*_1*c*_ = 1 s^−1^. The exchange matrix (Eq. 13) included *k*_*t*_ = 300 s^−1^ and *k*_*g*_ = 30 s^−1^ (see Fig. 2). These values were chosen to yield AXR values for *V* = −48 and *V* = −10 mV conditions (with *s*_o_ = 0) that are consistent with normal and ouabain-treated values observed experimentally [38].

DEXSY signals were simulated without noise and with parameters (timings, gradient, *b*_1_, *b*_2_, *t*_m_, etc.) set based on values used in the companion experimental study [38]. The low-field single-sided permanent magnet (PM-10 NMR MOUSE, Magritek) used in that study produced a *g* = 15.3 T/m static gradient in the active region, which was modulated with hard RF pulses for sub-millisecond diffusion encoding. Simulated timing parameters include (τ_1_, τ_2_) combinations (0.200, 0.735) ms and (0.593, 0.580) ms, which leads to (*b*_1_, *b*_2_) values (0.089,4.435) and (2.329, 2.179) ms/µm^2^ by Eq. 1 with *g* = 15.3 T/m.The *t*_m_ values were [0.2, 1, 2, 4, 7, 10, 20, 40, 80, 160, 300] ms. An example of the simulated DEXSY signals are shown in Fig. 3.

**Fig. 3:**
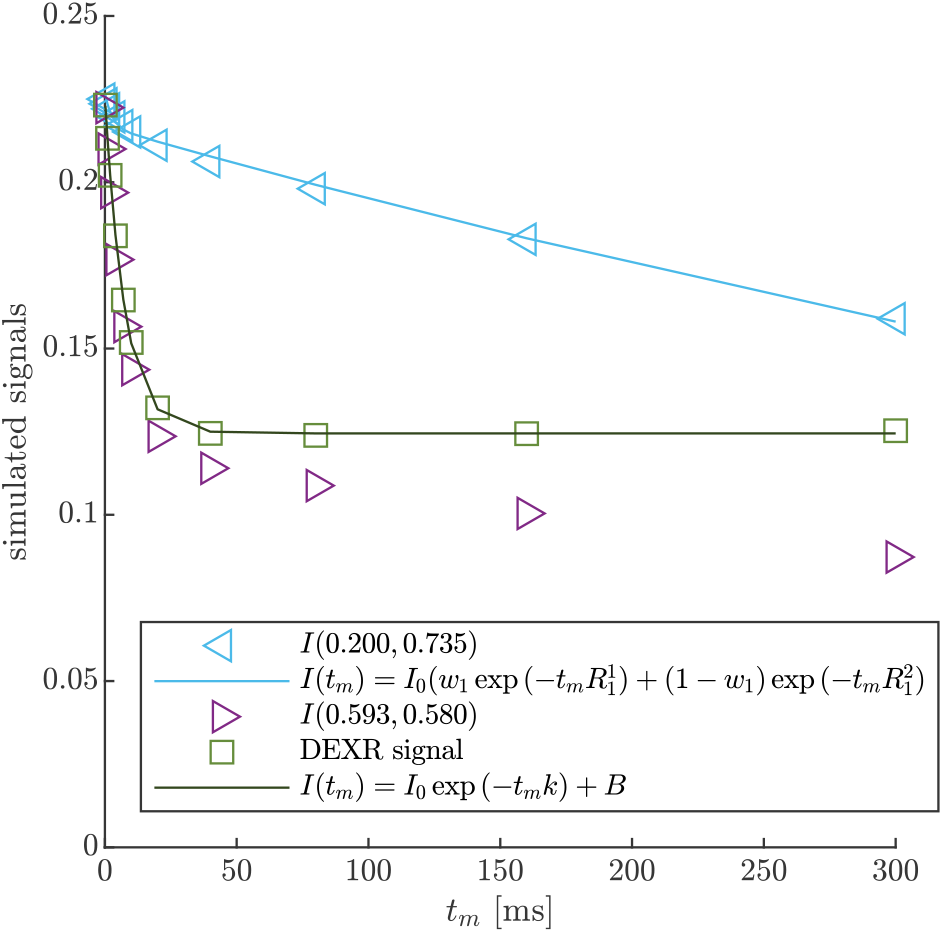
Example of DEXR method applied to simulated 3XM DEXSY data. Signals simulated for the normal media and *V* = −48 mV condition with ADC_*a*_ = 1, ADC_*b*_ = 1, and ADC_*c*_ = 0.1 µm^2^/ms using (τ_1_, τ_2_) combinations used to isolate exchange in the presence of *T*_1_ relaxation, i.e., the “DEXR signal”, and model fits. *I*(0.200, 0.735) decays primarily by *T*_1_ but with a slight initial decay due to some exchange weighting. This signal is fit with Eq. 20 (light blue solid line). *I*(0.539, 0.580) decays by exchange and *T*_1_. The *I*(0.539, 0.580) signal is divided by the Eq. 20 model fit to remove *T*_1_ relaxation and isolate exchange. The resulting DEXR signal is fit with Eq. 21 (solid black line). In this case, the estimated AXR = 132 s^−1^.

### 3.4. ADC *and* AXR *estimation*

ADC was predicted as the sum of the ADCs of each compartment multiplied by their volume fraction [127], e.g. ADC = *f*_*a*_ADC_*a*_ + *f*_*b*_ADC_*b*_ + *f*_*c*_ADC_*c*_ for the 3XM.

AXR was estimated using the DEXR method [90], following an approach similar to Method 3 in Ref. 83. First, to remove the effect of *R*_1_, the signal from (τ_1_, τ_2_) = (0.200, 0.735) was fit with a biexponential decay model,

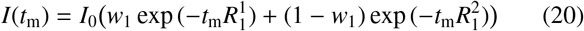

The resulting model was divided out of the signal from the (τ_1_, τ_2_) =(0.593, 0.580) experiment. The remaining “DEXR signal” was fit with a 3-parameter first-order rate model

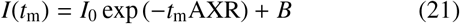

Note that while this was derived for a 2XM [83], here we use the model to fit numerical data simulated with both the 2XM and 3XM. In the case that the numerical data was simulated using a 2XM, the AXR estimated using Eq. 21 is expected to converge to the ground-truth *k* in Eq. 10, with a slight bias. In the case of the 3XM, the estimated AXR is weighted based on the ground-truth eigenvalues in Eq. 14. An example of the two fits involved in the DEXR method are shown in Fig. 3.

The slight bias noted above is due to the inability to acquire SGSE DEXSY signals with *b*_1_ = 0 since τ_1_ cannot be zero (Fig. 1). This slight diffusion weighting leads to some decay due to exchange in the (τ_1_, τ_2_) = (0.200, 0.735) point which then gets removed along with the *R*_1_ decay from the DEXR signal (see section 4.3.1 of Ref. [83]) This causes a bias in the estimated AXR (compare panels a–c to d–f in Supplementary Fig. S14 of Ref. [53]).

## 4. Results

### 4.1. Pump–leak model results

We will begin by using the PLM to simulate the effects of Na^+^/K^+^–ATPase inhibition and media ion and osmolyte perturbations on cell volume and voltage. The goal is to corroborate the results with experimental data from Ref. 38. Simulated cell volume changes are expected to be inversely related to experimentally-measured ADC changes (cell swelling leads to a greater fraction of restricted water and hence ADC reduction). The simulated voltages provide complementary insights, as voltage was not directly measured in these experiments.

Fig. 4 presents the steady-state volumes and voltages predicted by the PLM for various media conditions relative to that of the normal 128 mM NaCl, PUMP ON condition. The PUMP OFF condition models Na^+^/K^+^–ATPase inhibition by ouabain. The 128 mM NaCl, PUMP OFF condition results in an unstable state of gradual, continual swelling and depolarization due to an osmotic imbalance from intracellular impermeants. (See also Fig. 2 in Ref. 62.) We [38, 53] and others [128–130] observed ADC reduction in neural tissue after ouabain.

**Fig. 4:**
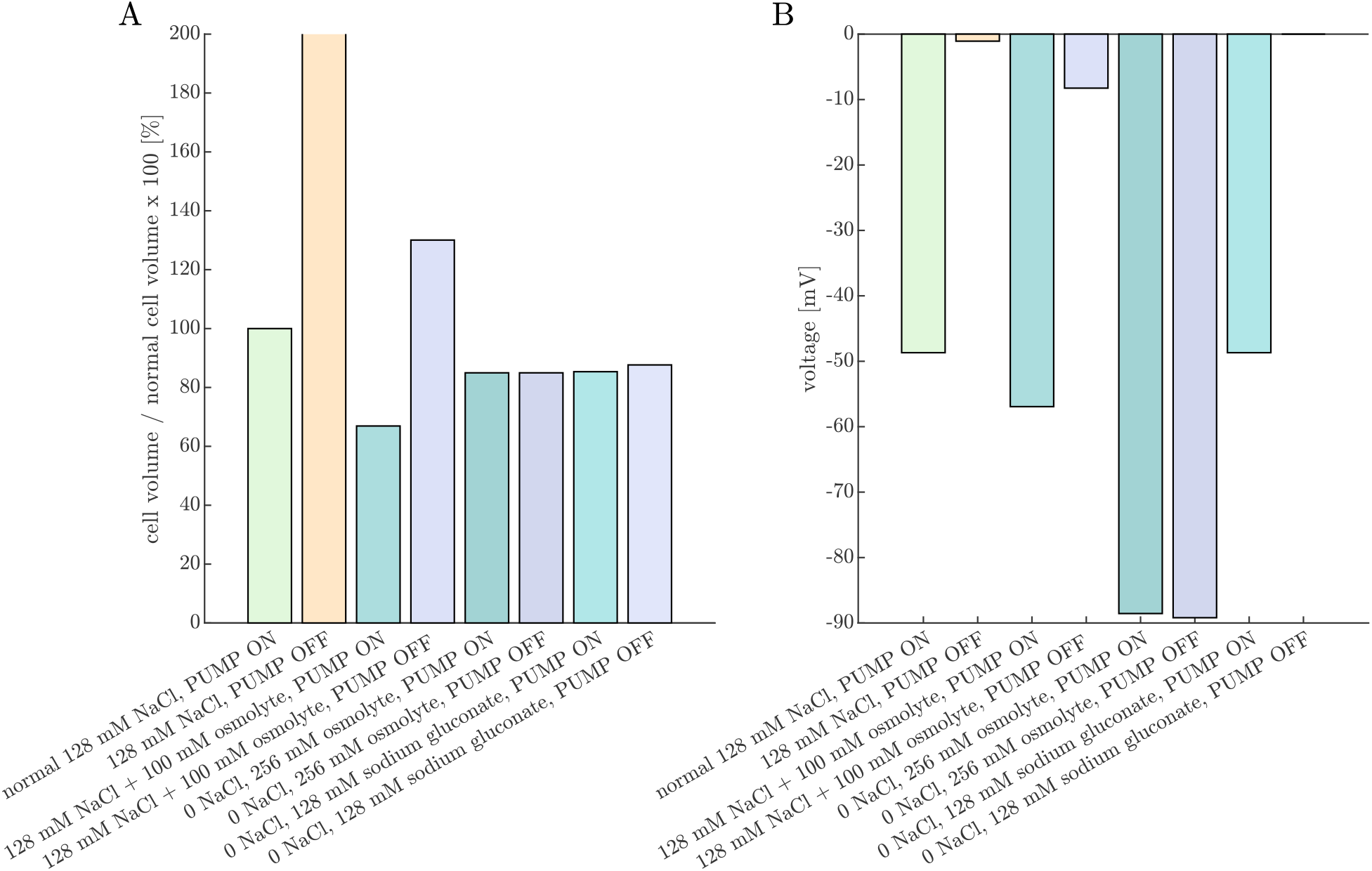
Pump-leak model (PLM) provides predictions of cell volume and voltage under conditions studied experimentally. Percent change in (A) cell volume and (B) voltage predicted under various Na^+^/K^+^–ATPase PUMP ON or OFF and media conditions after the PLM has run to steady-state (if applicable). With the 128 mM NaCl (and 4 mM KCl) media and PUMP OFF condition, the cell continues to swell and does not reach a steady-state. Addition of an osmolyte (+100 mOsm) reduces the volume and hyperpolarizes the cell in the PUMP ON condition, and stabilizes the volume but does not recover the voltage with the PUMP OFF. The cell volume is reduced to a similar level in both the PUMP ON and PUMP OFF conditions when NaCl is replaced by either sucrose or sodium gluconate (modeled as uncharged and monovalent anionic osmolytes in the bathing media, respectively). However, while voltage is similar between the PUMP ON and PUMP OFF conditions when NaCl is replaced by sucrose, full depolarization to *V* = 0 is predicted in the PUMP OFF condition with NaCl replaced by sodium gluconate. This figure complements experimental ADC and AXR measurements collected under similar conditions and presented in Figs. 2 and 3 of Ref. [38].

In an Aplysia CNS model, Jelescu *et al.* also observed an ADC decrease at the tissue level, but an ADC increase inside isolated neuronal soma [130]. This discrepancy may have resulted from soma being larger than the dephasing length (*l*_*s*_ > *l*_*g*_), leading to the localization regime. In the localization regime, cell swelling results in more water being further than *l*_*g*_ from plasma membranes, causing ADC to increase.

In contrast to the unstable swelling predicted by the PLM, we found [38, 53] that ADC eventually stabilized after ouabain administration, indicating a stable cell volume was reached. This is likely due to forces not accounted for in the PLM, such as the dura surrounding the spinal cord and trapped ECS osmolytes. This discrepancy was the inspiration for adding a fixed *w*_tot_ and *x*_o_ into the equation for the steady-state *f*_o_ (Eq. 19) which we use in the next section.

The PLM predicts cell shrinkage during the 128 mM NaCl + 100 mM osmolyte, PUMP ON condition, whereas experimentally we observed ADC to be only marginally affected by osmolytes under normal conditions [38]. This may arise because in reality the Na^+^/K^+^–ATPase pump rate may decrease under hypertonic conditions [131] whereas in the model the pump rate is constant. Additionally, the model does not include regulatory volume increase (RVI) mechanisms that recover cell volume in a hypertonic environment by increasing intracellular osmolarity [61]. The PLM predicts a stable but swollen volume during the 128 mM NaCl + 100 mM osmolyte, PUMP OFF condition, whereas we observed an increase in ADC above baseline, indicating cell shrinkage for ouabain-treated samples at 100 mOsm bath osmolarity. From Eq. 18, the osmolarity expected to recover volume is

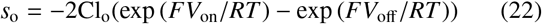

This equation is a function of the extracellular chloride concentration (Cl_o_) and voltage under the two conditions of PUMP ON and OFF (*V*_on_ and *V*_off_) and predicts volume recovery at *s*_o_ = 138 mOsm with Cl_o_ = 132 mM, *V*_on_ = −48 mV, and *V*_off_ = −10 mV. Therefore, the experimental results may differ from simulation predictions due to reduced *V*_on_, increased *V*_off_, or Cl_o_ being lower than the chloride concentration in the bath. The latter is plausible if repulsive forces from fixed negative charges of the extracellular matrix excludes some Cl_o_ from the ECS.

The PLM predicts that cells shrink when 128 mM NaCl is replaced by 256 mM of an uncharged osmolyte. The effect is similar under PUMP ON and OFF conditions. This is because Na^+^ and Cl^−^ are not fully confined to the ECS, so their osmotic contribution is less than their molar concentration. As a result, replacing 128 mM NaCl with 256 mM osmolyte creates a slightly hypertonic environment, leading to cell shrinkage. This aligns with experimental observations, where ADC increases slightly when switching to a 0 NaCl, 256 mM sucrose aCSF, with a similar effect observed when the Na^+^/K^+^–ATPase is inhibited by ouabain [38].

Replacing 128 mM NaCl with 128 mM sodium gluconate has a similar effect on cell volume. Since gluconate is a monovalent anionic osmolyte, it requires an equal concentration of cations to maintain electroneutrality. Thus, a 0 NaCl, 128 mM sodium gluconate solution has the same osmolarity as a 0 NaCl, 256 mM uncharged osmolyte solution. However, the PLM predicts a key difference between the two conditions in terms of membrane voltage under PUMP OFF conditions. In the 0 NaCl, 256 mM osmolyte solution, voltage remains unchanged between PUMP ON and PUMP OFF conditions because there is no Na^+^ to pump. In contrast, in the 0 NaCl, 128 mM sodium gluconate solution, the membrane voltage depolarizes from approximately –50 mV to 0 mV. This highlights why sodium gluconate is useful for testing whether certain mechanisms depend on voltage rather than volume [132].

In the companion experimental study [38], ADC and AXR were not affected when switching from normal aCSF to 0 NaCl, high sodium gluconate aCSF. From there, ADC and AXR increased when inhibiting the Na^+^/K^+^–ATPase with ouabain. This indicates that Na^+^/K^+^–ATPase was active prior to ouabain addition. Furthermore, this suggests that prior to ouabain addition, a subset of RVI mechanisms [61] that function with minimal chloride, utilizing Na^+^/K^+^–ATPase and downstream transport pathways, were functioning to maintain normal cell volume.

### 4.2. f_o_, two- and three-site exchange model results

With an understanding of how cell voltage and volume are connected to Na^+^/K^+^–ATPase activity and osmotic and ionic conditions, we now turn to the 2XM and 3XM. Both models employ the same analytical equation for the ECS fraction *f*_o_ (Eq. 19), so we start by looking at the dependence of *f*_o_ and osmolarities of extracellular and intracellular impermeants on voltage and osmolyte concentration *s*_o_ (Fig. 5).

**Fig. 5:**
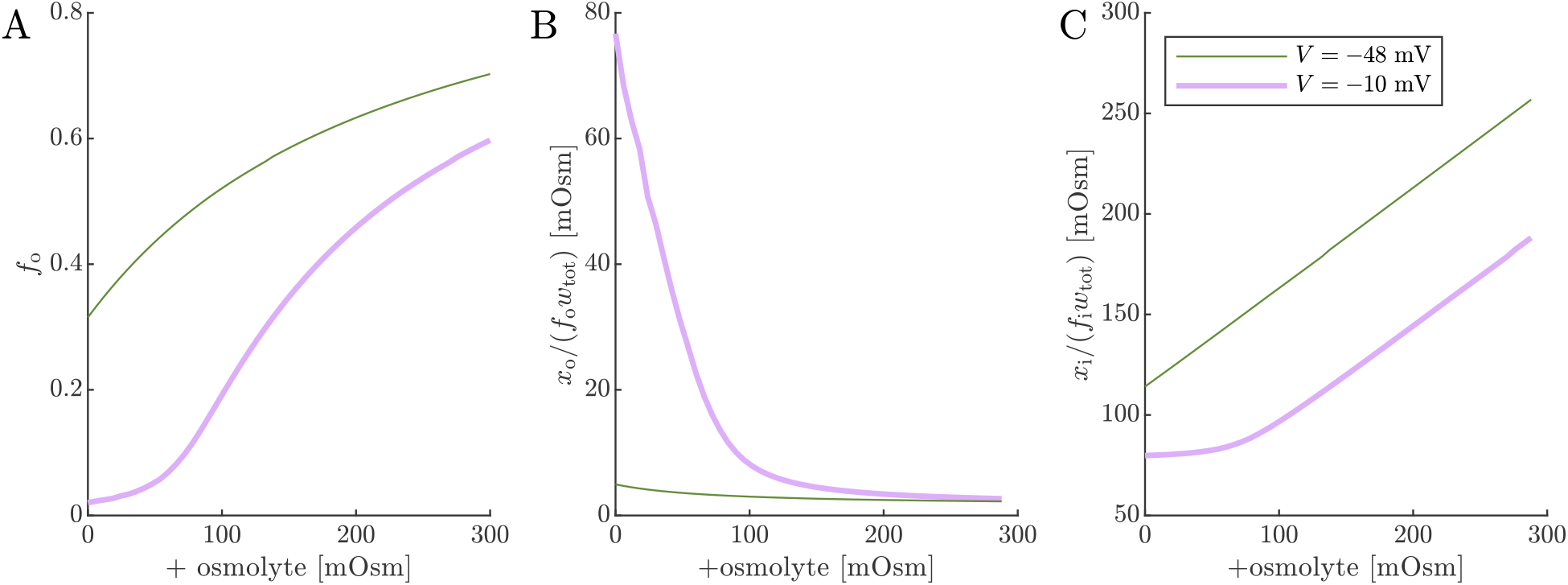
ECS volume fraction and osmolarities of extracellular and intracellular impermeants. Simulated (A) *f*_o_ from Eq. 19, (B) osmolarity of ECS impermeants, and (C) osmolarity of ICS impermeants for normal *V* = −48 mV and depolarized *V* = −10 mV conditions as a function of the osmolyte concentration *s*_o_. *f*_*a*_ is defined equal to *f*_o_ as the first step of numerical 2XM and 3XM simulations.

The effects of Na^+^/K^+^–ATPase activity and inhibition are modeled by setting *V* = −48 mV or −10 mV, respectively. *V* = −48 mV and *s*_o_ = 0 is considered the normal condition and leads to *f*_o_ = 0.32, slightly increased from the initial *f*_o_ = 0.3 due to the slight osmolarity of *x*_o_/ *f*_o_*w*_tot_. *V* = −10 mV and *s*_o_ = 0 is intended to model the effect of Na^+^/K^+^–ATPase inhibition. When *V* = −10 mV and *s*_o_ = 0, the fraction of chloride partitioning is reduced and *f*_o_ decreases to 0.02, *x*_o_/(*f*_o_*w*_tot_) increases substantially, and *x*_i_/(*f*_i_*w*_tot_) decreases. For the *V* = −48 mV case, increasing *s*_o_ from zero causes *f*_o_ to increase, but asymptotically due to the buildup of *x*_i_/(*f*_i_*w*_tot_). *x*_i_/(*f*_i_*w*_tot_) increases roughly linearly because there is very little effect from *x*_o_/(*f*_o_*w*_tot_). For the *V* = −10 mV case, *f*_o_ has a sigmoidal dependence on *s*_o_. At low *s*_o_, the dependence is shallow until it gradually overcomes *x*_o_/(*f*_o_*w*_tot_) and becomes steeper. At high *s*_o_, the dependence begins to level off as *x*_i_/(*f*_i_*w*_tot_) becomes more significant, similar to the behavior for the *V* = −48 mV case. When *x*_o_ is reduced, the plateau at low osmolarity becomes stronger, but at a value of *f*_o_ which is also reduced (data not shown). *f*_o_ = 0.02 at the plateau is roughly half the lower bound reported (as α) in real-time iontophoresis with tetramethylammonium studies of various *in vivo* and *ex vivo* CNS models involving ischemia, anoxia, or spreading depression/depolarization [123, 124]. This discrepancy could be because mechanical and hydrostatic pressures build up more strongly than predicted by the 1/ *f*_o_ or 1/ *f*_i_ scaling (as reported for ECM components [121]), contributing to the plateauing at both ends of the osmolarity range.

Next, results of ADC and AXR estimates from numerical 2XM simulations are shown (Fig. 6). ADC is simply a linear function of *f*_o_. The ground truth *k* = 300 s^−1^ is independent of *f*_o_. AXR estimates are biased slightly above *k* = 300 s^−1^, as discussed in Section 3.4, and the bias depends slightly on *f*_o_.However, relative to what we will see below, AXR estimates are not affected by osmotic condition. The 2XM is unable to explain the effect of ouabain and osmolytes on AXR which we observed experimentally [38].

**Fig. 6:**
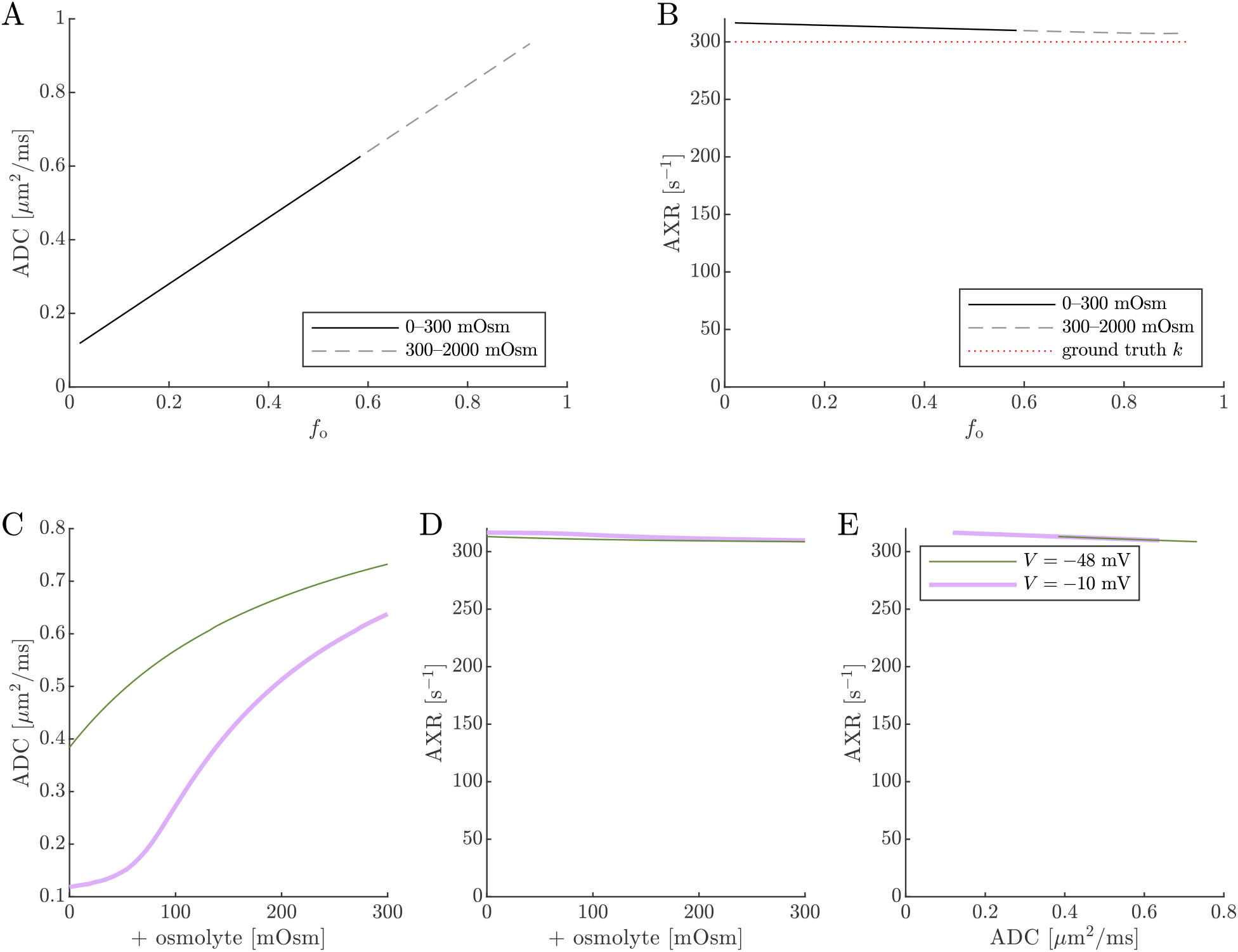
On data simulated using the 2XM, estimates of ADC are affected by osmotic condition, but estimates of AXR are not. A,B) Dependence of ADC and AXR on *f*_o_. The dashed lines show how the behavior extends as *f*_o_ increases towards 1. C,D) Dependence of ADC and AXR on osmolyte concentration *s*_o_ for the normal *V* = −48 mV and depolarized *V* = −10 mV conditions. E) correlation between ADC and AXR. In this 2XM with compartments *a* and *b*, ADC_*a*_ = 1 and ADC_*b*_ = 0.1 µm^2^/ms, and the ground truth *k* = 300 s^−1^ is independent of *f*_o_ or *s*_o_.

Next we show results of ADC and AXR estimates from numerical 3XM simulations (Fig. 7). As in the 2XM, the ADC is a function of *f*_o_ (Fig. 7 A). Unlike in the 2XM, the AXR is also a function of *f*_o_ (Fig. 7 B). When *f*_o_ is high, AXR approaches *k*_*t*_ = 300 s^−1^. When *f*_o_ is low, AXR approaches *k*_*g*_ = 30 s^−1^. In these extreme cases, the model effectively reduces to two-site exchange, and the agreement between the estimated AXR and the defined *k*_*t*_ and *k*_*g*_ serves as an internal validation. Inbetween, the AXR varies roughly linearly, with slight concavity consistent with (though not as pronounced as) a weighted sum of the eigenvalues.

**Fig. 7:**
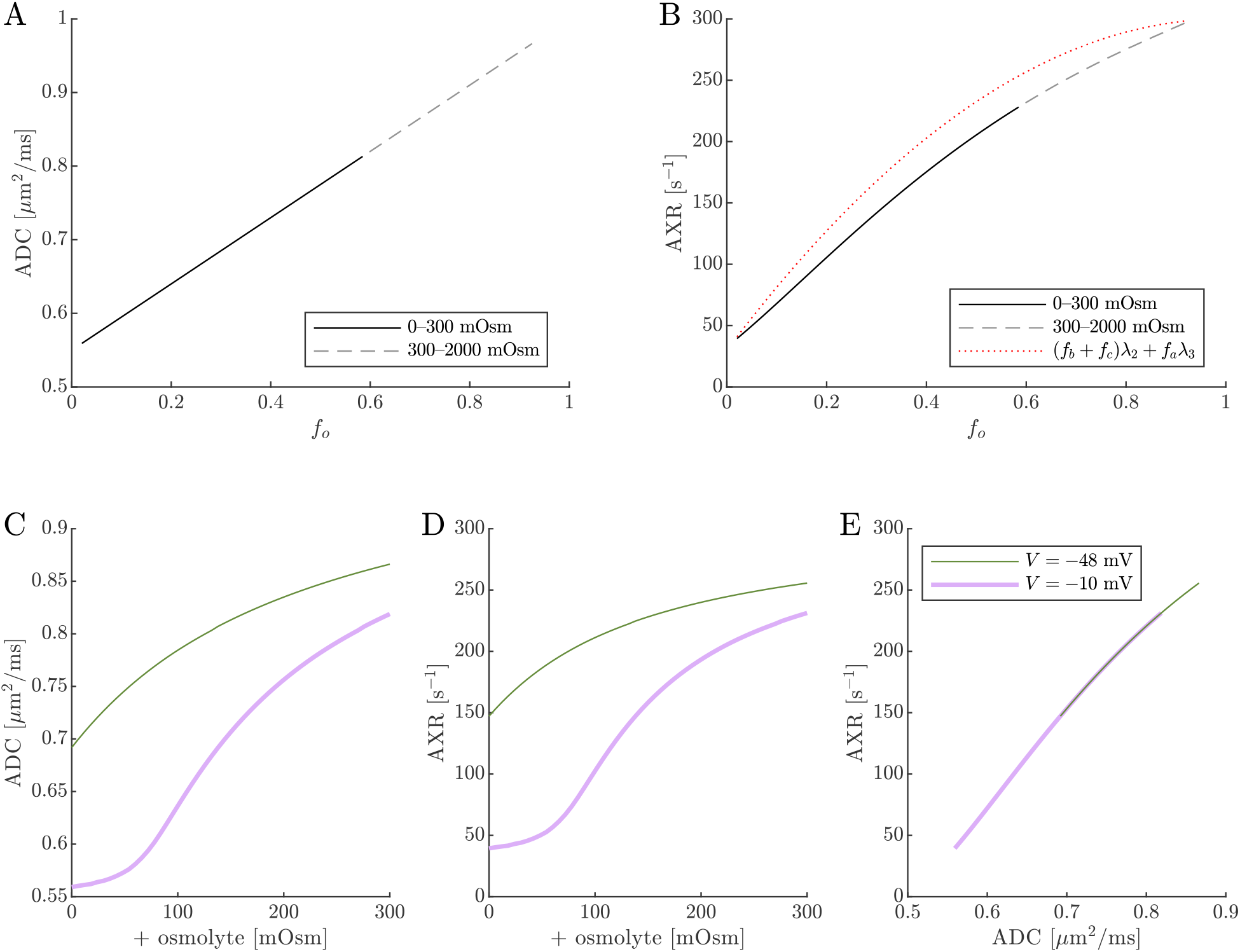
On data simulated using the 3XM, ADC and AXR are affected by osmotic conditions through a dependence on extracellular volume fraction. A,B) Dependence of ADC and AXR on *f*_o_. Exchange rates are expected to be a weighted sum of eigenvalues from Eq. 14 (dotted line). C,D) Dependence of ADC and AXR on osmolyte concentration *s*_o_ for the normal *V* = −48 mV and depolarized *V* = −10 mV conditions. E) correlation between ADC and AXR. Data was numerically simulated using the 3XM and ADC_*a*_ = 1, ADC_*b*_ = 1, and ADC_*c*_ = 0.1 µm^2^/ms.

The simulation predicts that transitioning from *V* = −48 mV to *V* = −10 mV causes AXR to drop from 140 s^−1^ to 40 s^−1^ and ADC to decrease by 23% (Fig. 7 C and D). Adding an osmolyte subsequently restores AXR and ADC. The trends appears somewhat sigmoidal due to their dependence on *f*_o_ and the sigmoidal relationship between *f*_o_ and osmolarity (compare Fig. 7 C and D to Fig. 5 A). This behavior was observed experimentally, though plateauing more completely and at concentrations above 100 mOsm (compare to Fig. 4 C and E in Ref. [38]). Hence the 3XM can qualitatively explain the effects of ouabain and osmolytes on AXR which we observed in the companion study [38]. The simulation predicts a roughly linear correlation between ADC and AXR, independent of voltage (Fig. 7 E), since ADC and AXR both depend on *f*_o_.

Diffusion exchange measurements are only sensitive to exchange between compartments with distinct water mobilities. Sensitivity increases as the compartment mobilities shift further apart. In the above simulation, ADC_*a*_ and ADC_*b*_ were set to 1 µm^2^/ms, making exchange between compartments *a* and *b* undetectable. This assumption—setting ECS and ICS diffusivities equal in axons aligned with *g*—is commonly used in the Neurite Orientation Dispersion and Density Imaging (NODDI) model [125], where it can introduce biases and degeneracy in parameter estimation [29]. Two plausible alternatives exist: (1) ADC_*b*_ > ADC_*a*_ or (2) ADC_*b*_ < ADC_*a*_. A number of studies on white matter favor the first scenario, concluding that intracellular diffusivity exceeds extracellular diffusivity [126]. To investigate this, we increased ADC_*b*_ to 1.5 µm^2^/ms (Fig. 8) and explored the opposite case by reducing ADC_*b*_ to 0.5 µm^2^/ms (Fig. 9). While varying compartmental ADC values affected the overall ADC, the magnitude of ADC changes, and the correlation between ADC and AXR, the qualitative behavior appears similar to the case that ADC_*b*_ = ADC_*a*_ (Fig. 7).

**Fig. 8:**
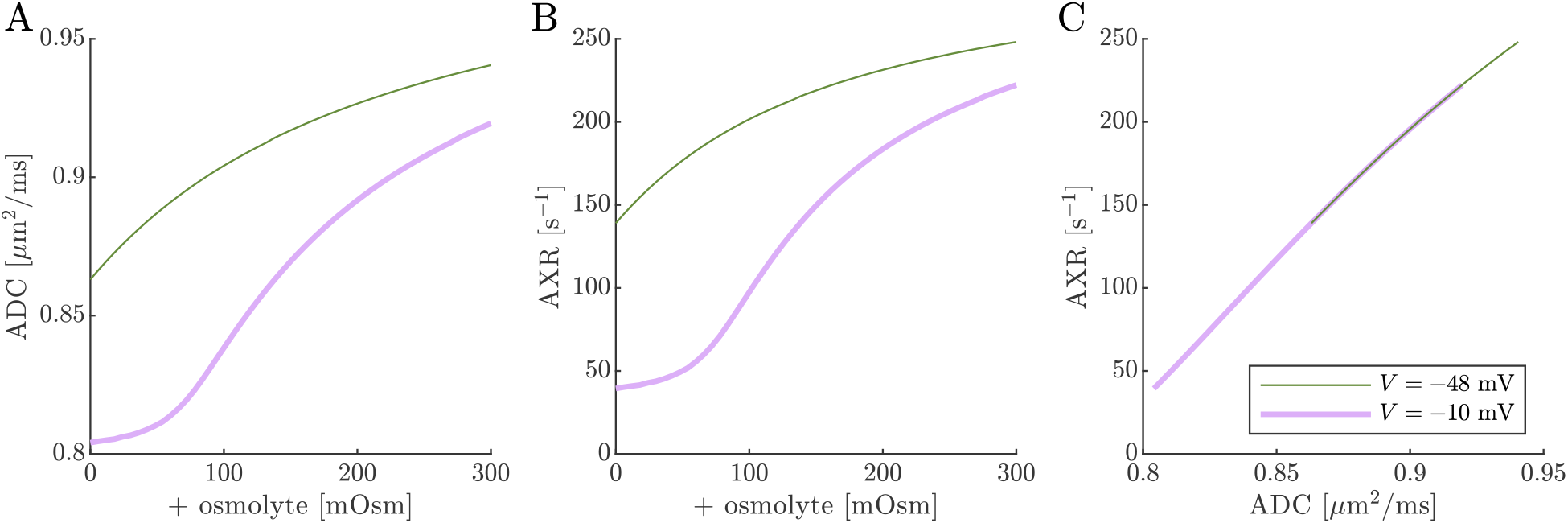
ADC and AXR estimates depend on ICS compartment mobility in the 3XM, ADC_*b*_ > ADC_*a*_ **case.** A,B,C) ADC, AXR, and the correlation between ADC and AXR predicted using the 3XM when an osmolyte is added to the normal media with *V* = −48 mV or −10 mV. In this model, ADC_*a*_ = 1, ADC_*b*_ = 1.5, and ADC_*c*_ = 0.1 µm^2^/ms.

**Fig. 9:**
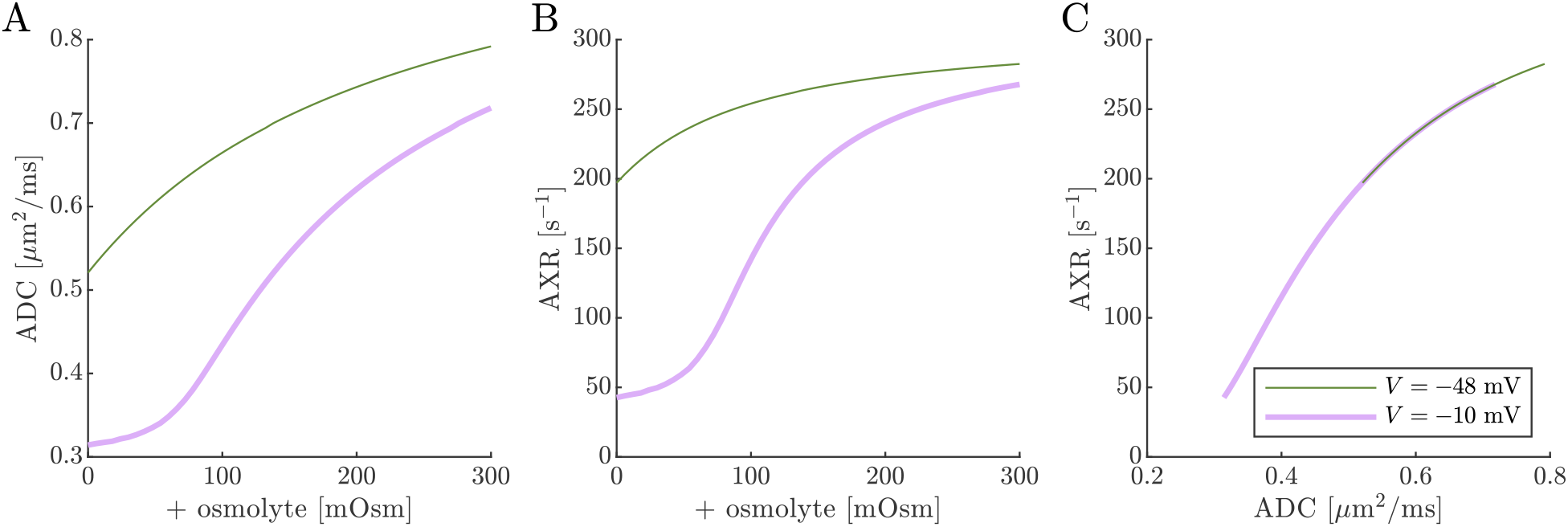
ADC and AXR estimates depend on ICS compartment mobility in the 3XM, ADC_*b*_ < ADC_*a*_ **case.** A,B,C) ADC, AXR, and the correlation between ADC and AXR predicted using the 3XM when an osmolyte is added to the normal media with *V* = −48 mV or −10 mV. In this model, ADC_*a*_ = 1, ADC_*b*_ = 0.5, and ADC_*c*_ = 0.1 µm^2^/ms.

Cell swelling and reduction of *f*_o_ is expected to reduce the diffusivity in the ECS [133]. Diffusion models such as NODDI account for this by having the ECS diffusivity depend linearly on *f*_o_ [29, 125]. Following them, in Fig. 10 we set ADC_*a*_ =1.7 *f*_o_. Doing so leads to a non-monotonic relationship for ADC when *V* = −10 mV, affecting the correlation between ADC and AXR. However, the behavior for AXR appears similar to previous cases.

**Fig. 10:**
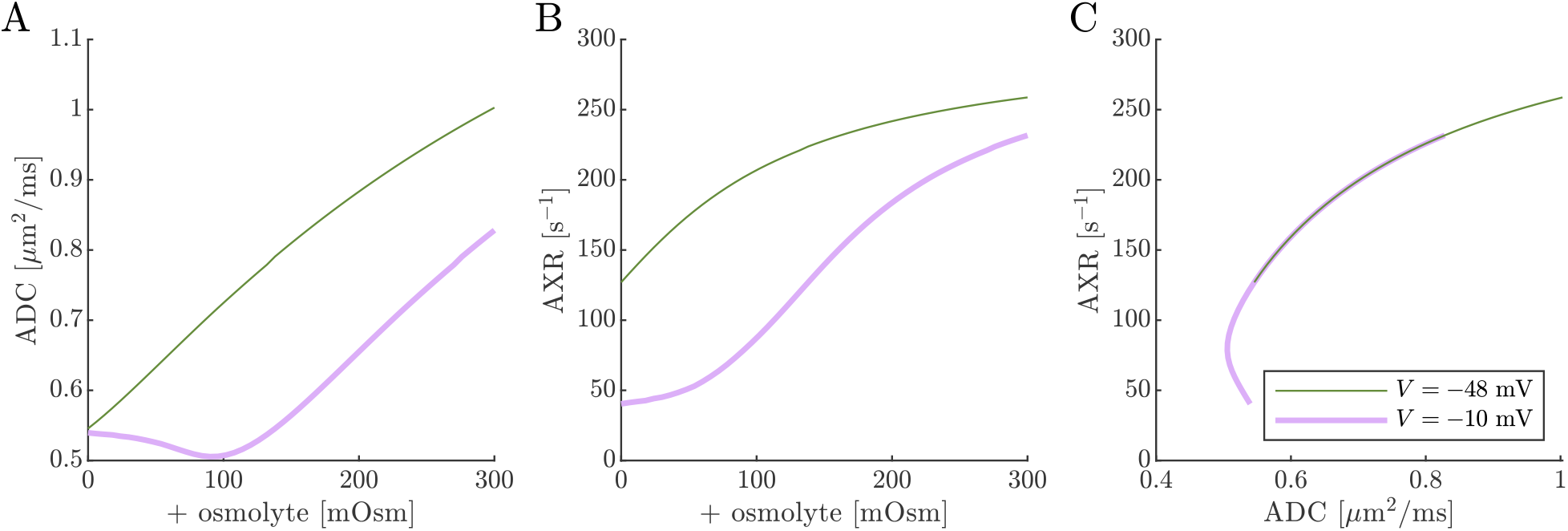
ADC and AXR estimates depend on ECS compartment mobility in the 3XM. A,B,C) ADC, AXR, and the correlation between ADC and AXR predicted using the 3XM when an osmolyte is added to the normal media with *V* = −48 mV or −10 mV. In this model, ADC_*a*_ = 1.7 *f*_o_, ADC_*b*_ = 1, and ADC_*c*_ = 0.1 µm^2^/ms.

While the 3XM is able to predict the major effects of ouabain and osmolytes on AXR, it does not fully capture our experimental findings [38]. In particular, the model predicts osmolytes to increase AXR when *V* = −48 mV and cannot capture our experimental findings that AXR is unaffected by addition of up to 30 mOsm and only marginally affected at higher concentrations while the Na^+^/K^+^–ATPase is active [38]. Additionally, since the model predicts the correlation between AXR and ADC with osmotic treatment to be roughly linear and independent of Na^+^/K^+^–ATPase activity, it does not explain why ADC and AXR can change independently, as we observed during oxygen and glucose deprivation studies (Fig. 4 in Ref. [53]). Furthermore, the model does not capture the sigmoidal correlation between AXR and ADC under ouabain treatment or the relatively flat and distinct correlation under normal conditions, both of which were observed experimentally (Fig. 4F in [38]). These discrepancies suggest the presence of additional mechanisms beyond those included in the model. Possibilities include the role of volume regulation [61], osmotically-induced lipid phase transitions [134, 135], overall tissue volume changes [136, 137], cell shape changes [138], neurons and glia acting differently [139, 140], and pressures not accounted for in the model [121, 141]. Other potential contributing factors are explored below.

Previous studies on *ex vivo* neonatal mouse spinal cord have shown that the DEXR signal has multiexponential character (see Fig. 4.7 in Ref. [142]). This could arise from the system containing multiple exchange processes with different rate constants, i.e., an exchange rate constant distribution [37]. Here we tested whether the 3XM could result in multiexponential character. Fig. 11 shows simulated signals for 100 mixing times spaced on a log_10_ scale from 0.2 to 400 ms for conditions defined by (*V, s*_o_) =(−10 mV, 0 mOsm), (−48 mV, 0 mOsm) and (−48 mV, 100 mOsm). In all cases, the signal was well-fit by a single AXR (Eq. 21). The behavior expected based on the two eigenvalues also appears roughly monoexponential because the eigenvalues are not too dissimilar. Hence, the 3XM alone can-not explain the stark multiexponential character of the DEXR signal observed experimentally and suggests that either the exchange process deviates from first-order kinetics [39] or the tissue contains multiple microenvironments which are each characterized by their own AXR.

**Fig. 11:**
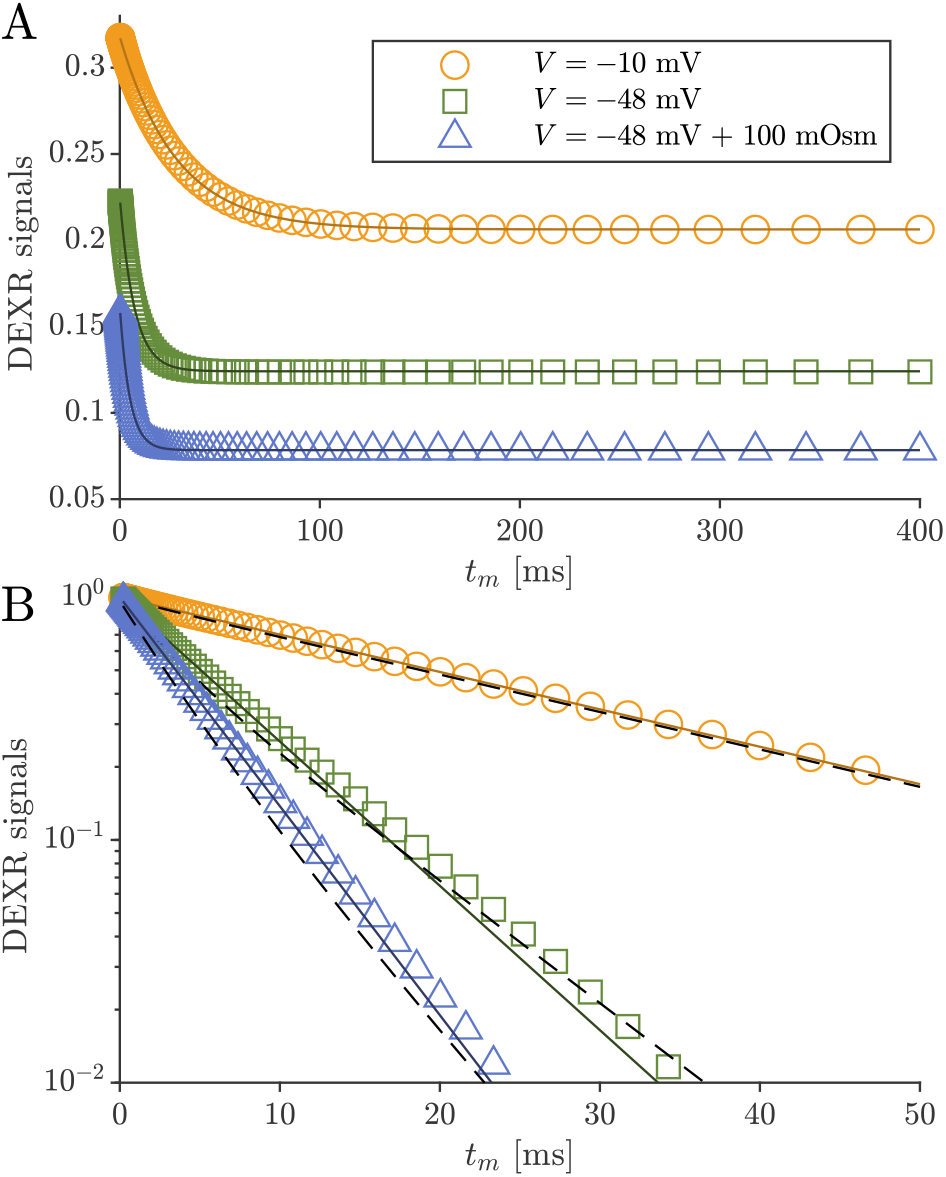
Simulated data and model fits show roughly single exponential decay. DEXR signals simulated for depolarized *V* = −10 mV, normal *V* = −48 mV and normal *V* = −48 mV + 100 mOsm conditions and with ADC_*a*_ = 1, ADC_*b*_ = 1, and ADC_*c*_ = 0.1 µm^2^/ms along with AXR fits (Eq. 21) showing the full decay on a linear scale and (B) the initial decay on a semi-log scale.In (B), biexponential models with the form *I*(*t*_m_) = (*f*_*b*_ + *f*_*c*_) exp(−λ_2_*t*_m_) + *f*_*a*_ exp(−λ_3_*t*_m_) with eigenvalues defined by Eq. 14 are also shown (dashed lines). For the three conditions λ_2_ = 35.5, 115, and 171 s^−1^ respectively, and λ_3_ is always 300 s^−1^.

## 5. Discussion

We have developed a 3XM for DEXSY of CNS gray matter tissue, incorporating passive transmembrane and geometric water exchange between an extracellular compartment and two intracellular compartments. This model establishes a fundamental mechanism by which AXR varies with osmotic conditions, as it is directly influenced by *f*_o_. As the ECS fraction increases, AXR is biased toward the faster *k*_*t*_, whereas with less ECS, AXR is dominated by the slower *k*_*g*_.

The simple behavior of *f*_o_ predicted by Eq. 19 and its effect on the 3XM also explains why AXR does not depend on Na^+^/K^+^ pump activity *per se*. Under normal conditions, ions are partitioned by Na^+^/K^+^ pump activity. This maintains normal *f*_o_ and AXR. Inhibiting the Na^+^/K^+^ pump causes ions to redistribute to their Donnan equilibrium and cell swelling/ECS shrinkage, reducing AXR through its dependence on *f*_o_ and not on Na^+^/K^+^ pump activity *per se*. Adding osmolytes or substituting NaCl with osmolytes restores normal cell volume and *f*_o_, seemingly rescuing AXR without restoring Na^+^/K^+^ pump activity. This behavior cannot be explained by the 2XM where *k* is independent of *f*_o_. This behavior also cannot be explained by a 2XM where the AXR is a sum of active and passive exchange rate constants [48, 50, 53].

### 5.1. Limitations and future work

The DEXR method was originally developed to measure two-site exchange [89, 90]. First, the method involves two measurements at a constant *b*_1_ +*b*_2_ which is chosen to maximally attenuate spins in the more mobile compartment while also maximizing coherence of spins in the less mobile compartment. While multiple sites can be resolved with more *b*_1_ + *b*_2_ combinations, this comes at a cost of increased experimental time [89]. Similar assumptions and tradeoffs apply to FEXSY and FEXI [91, 92]. The feasibility of using FEXI to measure different exchange processes in human brain by varying *b*_1_ for the diffusion filter has been demonstrated, but still assuming each of those processes follows two-site exchange [34]. Even more deeply ingrained in the method, the DEXSY pulse sequence with two diffusion encodings separated by a mixing time is considered the best sequence for measuring exchange between two compartments [143], but is not necessarily the best for measuring exchange among three or more compartments. Better methods more amenable to revealing and estimating parameters of a multisite model should be developed going forward.

That said, the state of the field and the challenges should be noted. First, multisite exchange is much more developed for REXSY than for DEXSY [96, 103–105, 144]. This is because the CPMG train of the second *T*_2_ encoding block doubles as a signal readout so that more data points can be acquired in less time. Additionally, relaxation follows a predictable exponential kernel in the motional narrowing regime [145]. With more data points and a predictable kernel the 2-D inversion can be performed to reveal multiple sites and their connectivity [2, 5, 86, 97, 146–150]. Adaptation of DEXSY-based methods for multisite exchange is hampered by the time required to acquire individual *b*_1_, *b*_2_ combinations and the absence of a unified kernel for free and restricted components. Efforts to reduce the required data with constraints [87, 88], ultrafast Laplace methods [3, 151], or modulated gradient spin echo (MGSE)– based diffusion signal readouts [152–154] show promise, but incorporating mixed kernels [155] is challenging because the decay profile (e.g. motional averaging vs. localization) depends on the size of the restriction which is not known *a priori* [72].

The model calculates *f*_o_ assuming equal osmolarities in the ECS and ICS and does not account for a recently proposed water barochemical pressure gradient [141]. Compartment mobilities are modeled as ADC values, assuming there to be a distinct and fixed number of compartments, ignoring the possibility of non-Gaussian diffusion and the true nature of tissue heterogeneity. SVR changes and their effect on AXR are not modeled directly. We model the cell as being isolated, not accounting for intercellular exchange which could be significant with *k*_*t*_ > *k*_*g*_. We ignore microstructural changes such as neurite beading which occur during Na^+^/K^+^–ATPase inhibition [156] and affect signal differently depending on the encoding time [157]. Exchange is modeled as a first-order rate processes, which is only valid for barrier-limited transmembrane exchange [27, 90] and not for geometric exchange [39]. For instance, spins near points of branching will tend to exchange faster than spins further away along the same process. All of these factors could contribute to the discrepancy between 3XM predictions and experimental findings. These effects could be accounted for in future Bloch–Torrey [77, 158, 159] or random walk models that utilize more realistic branching geometries [42, 160, 161] rather than compartments, or by incorporating cell type-specific [24, 35] or geometry-specific motional averaging models [74] with anisotropy [162].

The hope of this study is that future diffusion modeling in gray matter will consider the importance of multisite exchange at PGSE MRI-relevant time and length scales. While additional validation is needed, the sensitivity of AXR measurements to transmembrane and geometric exchange pathways could provide an exciting direction for its development as an MRI biomarker.

## 6. Conclusion

NMR is uniquely suited for quantifying steady-state exchange and has a long track record in this area. However, its broader impact has been limited by the fact that determining exchange rate constants is an inverse problem requiring modelbased interpretation. Most methods, including DEXR, impose a 2XM, which can confound interpretation when applied to systems with more than two exchanging components. We propose that gray matter is such a system—one that exhibits both transmembrane and geometric exchange between one ECS compartment and two ICS compartments.

In this theoretical study, we investigated how applying a 2XM-based method to simulated 3XM DEXY data affects AXR measurements. Our simulations show that changes in osmotic and ionic conditions alter compartmental volume fractions, which in turn influence the AXR—behavior not predicted by the 2XM.

We found that the theoretical AXR and ADC predictions qualitatively match experimental observations reported in a companion study [38] across a wide range of osmotic conditions. These results offer an alternative explanation to hypotheses involving active water cycling [48, 50, 53] and represent a first step toward incorporating multisite exchange into biophysical modeling of microstructure in gray matter [126].

## Data and code availability statement

MATLAB code files to perform simulations and generate figures in this paper have been made publicly available through the following GitHub repository: https://github.com/nathanwilliamson/MultisiteExchange

## CRediT author statement

**Nathan Williamson:** Conceptualization, Theory, Methodology, Investigation, Analysis, Writing. **Rea Ravin** Conceptualization, Theory, Methodology, Investigation, Writing. **Teddy Cai:** Conceptualization, Theory, Methodology, Writing. **Julian Rey:** Conceptualization, Theory, Methodology, Writing. **Peter Basser:** Conceptualization, Theory, Methodology, Writing.

## Declaration of competing interests

The authors have no conflicts of interest to disclose. The views, information or content, and conclusions presented do not necessarily represent the official position or policy of, nor should any official endorsement be inferred on the part of, the Uniformed Services University, the Department of Defense, the U.S. Government or the Henry M. Jackson Foundation for the Advancement of Military Medicine, Inc.

### Declaration of generative AI and AI-assisted technologies in the writing process

During the preparation of this work the author(s) used Chat-GPT in order to edit for grammar and clarity. After using this tool/service, the authors reviewed and edited the content as needed and take full responsibility for the content of the published article.

## Acknowledgments

This work was partially funded by the Department of Defense in the Military Traumatic Brain Injury Initiative (MTBI^2^) under award HU0001-24-2-0051. NHW, RR, TXC, and PJB were also supported by the IRP of the NICHD, NIH. JAR was supported by an NIGMS Postdoctoral Research Associate Training (PRAT) Program Fellowship Award (Project Numbers: 1FI2GM150429-01; 1ZIEGM000002-18).

## Notes

https://github.com/nathanwilliamson/MultisiteExchange

### Competing Interest Statement

The authors have declared no competing interest.

### Summary of Updates

This version corrected an error in Eq. 4

